# Determinants of membrane sensitivity to the peptide MP-1 (*Polybia paulista*)

**DOI:** 10.1101/2025.09.17.676910

**Authors:** Andrew Booth, Arindam Pramanik, Dagmara Kobza, Didi Derks, Lekshmi Kailas, William J. Brackenbury, Simon D. Connell, Thomas A. Hughes, Paul A. Beales

**Affiliations:** School of Chemistry and Astbury Centre for Structural Molecular Biology, University of Leeds, Leeds, UK; School of Medicine, Wellcome Trust Brenner Building, St James’s University Hospital, University of Leeds, Leeds, UK; Amity Institute of Biotechnology, Amity University, Delhi, India; School of Physics and Astronomy and Astbury Centre for Structural Molecular Biology, University of Leeds, Leeds, UK; York Biomedical Research Institute, Department of Biology, University of York, Heslington, York, UK; School of Science, Technology and Health, York St. John University, Lord Mayor’s Walk, York, UK

## Abstract

The membrane-disrupting peptide Mastoparan-1 (MP-1), derived from the wasp *Polybia paulista* is known to possess antimicrobial properties, and exhibits enhanced activity against a number of cancer cell lines relative to healthy cells. Due to the mechanism of action of MP-1 it is likely that differences in plasma membrane lipid composition arising from cancer associated mutations, such as localisation of phosphatidylserine (PS) lipids to the outer leaflet of the plasma membrane, are involved in driving that enhanced activity. Rapid screening of MP-1 mutants in a combined approach using model membrane and cell-based biological assays, has led to the identification of a number of derivative peptides with enhanced selectivity for cancer-like membrane models and breast cancer cell lines and provided insights into the mechanism of membrane disruption and cell death. Notably, the morphology of the membrane perturbations observed by Atomic Force Microscopy (AFM) and activity in cell model systems can change considerably in response to single-point mutations in the MP-1 sequence, indicating a complex structure-activity relationship.

## Introduction

Mastoparan-1 (‘MP-1’, [IDWKKLLDAAKQIL-NH_2_]) is an amphiphilic peptide derived from the wasp *Polybia paulista*^1^. MP-1 has been studied for its antimicrobial activity^2^, and has exhibited enhanced cytotoxicity against cancer cell lines including prostate and bladder cancer^3,4^, without inducing haemolysis^1^. Membrane-disrupting peptides such as MP-1 cause lysis of the cell membrane leading to cell death^5^. The enhanced activity of these peptides against certain cancers is driven, at least in part, by differences in plasma membrane lipid composition in cancer cells resulting from disrupted membrane homeostasis^6,7^.

The membrane selectivity of MP-1 has been shown to be enhanced by the presence of cholesterol^8^ and phosphatidylserine (PS) lipids, with synergistic enhancement when PS and phosphatidylethanolamine (PE) lipids are both present^9^. Many cancers have increased exposure of PS and PE headgroup lipids on the outer leaflet of the plasma membrane due to reduced activity of the flippase enzymes that ordinarily maintain the location of these lipids on the cytosolic leaflet, as also observed in early apoptosis^7,10,11^. The influence of charged headgroup lipids on MP-1 activity can be rationalised by enhanced electrostatic association between the positively charged MP-1 molecule and a negatively charged membrane. Whereas the distinct behaviour observed in membranes containing multiple phases, or lipids that influence local membrane curvature, suggests that selectivity may also rely on the ability of the peptide to locally influence lipid self-assembly (e.g.: phase behaviour, curvature)^8^.

Limited prior studies have investigated a small number of MP-1 mutants, notably with regards to its capacity to induce degranulation in mast cells^12^, and membrane disruption in model membrane vesicles^13^. Ye *et al*. identified key features, such as amidation of the C-terminus and the integrity of the hydrophobic face as critical for its activity in mast cells^12^. While Leite *et al*. found that replacement of the aspartic acid residue in position D2 with asparagine had a significant effect on the conformation of the peptide in contact with phospholipid membranes and a concurrent diminishing of disruption in vesicle membranes^13^. Both studies point to a more subtle relationship between the amino acid sequence and the membrane activity of MP-1, particularly implicating conformational properties over bulk physiochemical characteristics.

With this in mind, we determined to use a combination of biological and biophysical assays to screen variants of MP-1 and identify those with improved characteristics as candidate anti-cancer agents. By initially producing a subset of single residue substitution variants, we hoped to identify key residues involved in membrane specificity and overall membraneolytic potential. Rapid screening of mutants was facilitated by the large unilamellar (LUV) fluorescent dye leakage assay, which is well-established in this role. Biophysical assays including atomic force microscopy (AFM) of supported lipid bilayers (SLBs) were then undertaken to investigate the mechanism of membrane disruption in model membranes and to identify changes in mechanism. Initially, simplified membrane compositions of DOPC and 80:20 DOPC:DOPS have been employed as model non-cancer and cancer membranes, respectively. Finally, candidate mutants were taken forward into biological assays with ‘cancer’ and ‘non-cancer’ model cell lines to assess how improvement in selectivity and potency in model membrane systems relates to behaviour in cell models. While the membrane activity of these peptides is the basis of their activity, model membrane compositions will not precisely match cell plasma membrane or cellular characteristics such as membrane repair processes, which vary by cell type^14^. Further variants of MP-1 can then be produced based on these findings, leading to refined candidate peptides.

## Results and Discussion

### Cell lines demonstrate a wide range of sensitivities to MP-1, but cancer cells are not significantly more sensitive

Our first aim was to determine the sensitivity of a range of mammalian cell lines to MP-1 in an effort to narrow down characteristics that define sensitivity. A range of lines derived from either human cancers or from non-transformed origins were treated with different doses of MP-1 and survival was assessed relative to untreated using MTT assays (**Fig 1A** and **B**). IC_50_ values varied almost 6-fold from most sensitive (19 μM, breast cancer line AU565) to most resistant (111 μM, non-transformed breast epithelial line MCF10A). Cancer cell lines (**Fig 1A**) were not significantly more sensitive than non-cancer lines (**Fig 1B**) (T Test p>0.05), demonstrating the reported cancer-specificity of MP-1 may be an over-simplification.

**Figure 1.**
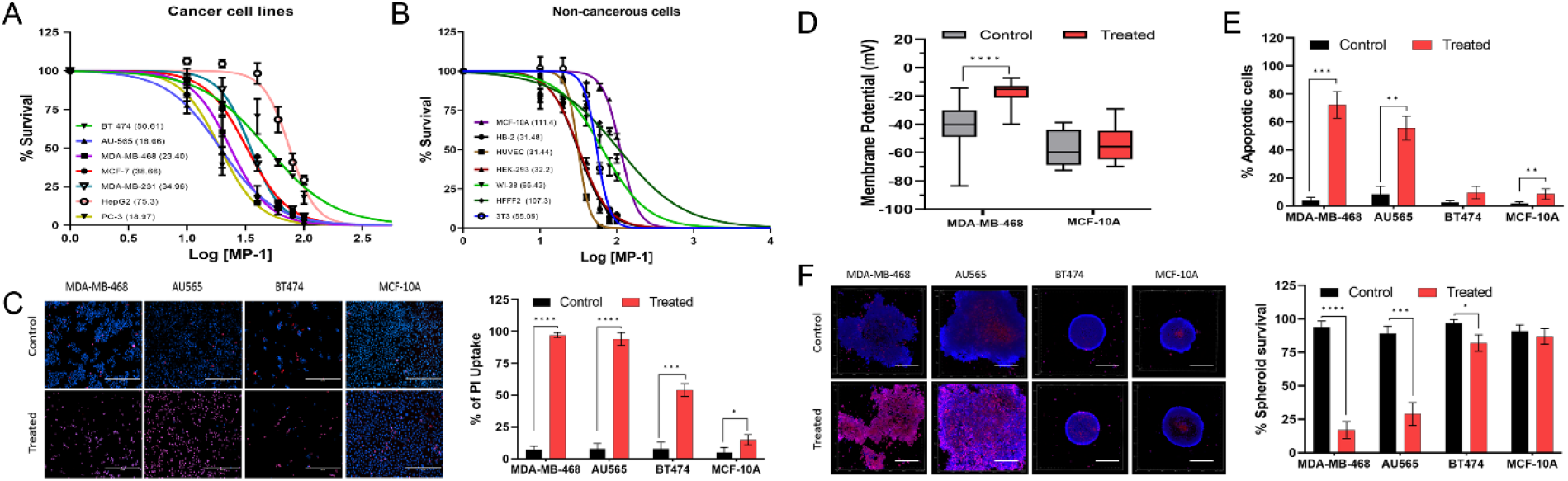
Cytotoxicity and membrane perturbation of cancerous and non-cancerous cell lines by MP1. MTT assays of cell viability after treatment with 0 – 200 µM concentrations of MP-1 in **A**: Cancer cell lines and **B**: Non-cancerous cells. The IC_50_ values are denoted in the figure for respective cell lines in µM. **C**: Propidium iodide membrane permeability assay, fluorescence microscopy of cells treated with an IC_50_ dose of MP-1 and untreated control cells. (Scale bar 200 µm). **D**: Whole cell patch clamp assay, showing membrane potentials for MDA-MB-468 (cancer-like) and MCF-10A (non-cancer-like) treated with IC_50_ dose of MP-1 and untreated controls. **E**: Flow cytometry apoptosis assay. Percentage of apoptotic cells determined by Annexin V-FITC and 7-AAD staining, for cells treated with IC_75_ MP-1 (30µM, 35µM, 70µM and 140µM for AU565, MDA-MB-468, BT474 and MCF10A respectively) or untreated controls. **F**: 3D spheroid assay. Spheroids treated with IC_50_ of MP-1 for 24 h followed by staining with Hoechst 33342 and propidium iodide to determine % survival of cells using confocal fluorescence microscopy (Scale bar 200µm).

Next, we investigated the characteristics of the growth inhibition/cell death induced by MP-1. For further study, we selected 4 representative cell lines, all from the same tissue of origin (breast epithelium) and covering a range of sensitivities (the most sensitive, AU565; the least sensitive, MCF10A; two intermediate, MDA-MB-468, BT474). We treated these four lines with their IC_50_ doses of MP-1, as defined above, and assessed plasma membrane permeability using a pair of DNA-binding fluorescent dyes: propidium iodide, which is not cell-permeable therefore can only access cell nuclei if membrane integrity is lost, and Hoechst 33342, which stains all cell nuclei (**Fig 1C**). All cell lines showed significant increases in proportions of propidium iodide positive cells after treatment. However, the more resistant cells showed smaller increases, even though these assessments were made using the IC_50_ doses appropriate for each cell line, hinting at differences in mechanism of action between relatively sensitive and resistant lines. Furthermore, we demonstrated by patch-clamping that treatment with the IC_50_ dose of MP-1 in the relatively sensitive MDA-MB-468 is associated with a loss of plasma membrane potential within 10 min of MP1 addition, while this was not the case in the resistant MCF10A (**Fig 1D**). To investigate contributions of apoptotic death to these responses, we performed annexin V/7-AAD staining in these cells, either untreated or treated with IC_75_ doses of MP-1 (**Fig 1E; Fig S1** for representative dot plots). The relatively resistant cell lines showed very low levels of MP-1 dependent apoptosis (BT474, MCF10A ; both <10%), as compared to the more sensitive lines that were up to 70% apoptotic (AU565, MDA-MB-468). None of the cell lines showed MP-1 dependent depolarization of the mitochondrial membrane (**Fig S2**). Finally, we assessed the responses to MP-1 of these cell lines in 3D spheroid cultures, using the fluorescent dyes as above (**Fig 1F**). We found the sensitive cells (AU565, MDA-MB-468) showed substantial cell lysis, while resistant lines demonstrated little lysis in this context. We concluded that MP-1 acts on different cell types with a range of efficacies, but without cancer-specificity, and that it potentially acts through more than one mechanism. Furthermore, the primary mode of action of MP-1 againsts sensitive cancer cell lines is consistent with plasma membrane damage, as has previously been reported due to cell uptake of membrane impermeable dyes within minutes of MP-1 addition and visualisation of direct damage^4^. Here, we observe loss of membrane potential within 10 min. While apoptosis leads to loss of membrane integrity, this would not be observed until ∼12 h^15^ and is therefore downstream of the direct membrane damage caused by MP-1.

### Individual residues within MP-1 and membrane constituents define MP-1 activity in membranes

Our next aim was to investigate the functional role of individual residues within MP-1 with respect to membranolysis. We employed an *in vitro* assay measuring release of a fluorophore from synthetic membrane vesicles which is suitable for high throughput screening, in order to determine which residues or residue properties are required for MP-1 function and thereby design mutants that might have improved cancer specificity and/or potency.

An initial set of MP-1 mutants were selected with the sequences shown in **Supplementary Figure 1**. Of these, twelve comprised an alanine scan to identify key residues, while Lysine or Histidine substitutions were made to examine the role of charge, and a single substitution for a glutamine residue was made to probe the role of peptide-peptide interactions. As MP1 is known to form an amphipathic helix upon interacting with lipid membranes, the helical wheel structure of the mutant peptide was considered in their design (see SI **Figure S7**).

An automated assay was developed to rapidly screen MP-1 mutants based on their ability to disrupt model membranes of various compositions relative to unmodified MP-1. The presence of PS headgroup lipids on the outer leaflet of certain cancers has been implicated as a possible factor in the enhanced activity of MP-1 against these cells^9^. Therefore, parallel leakage assays were performed with vesicles composed of 80:20 DOPC:DOPS and DOPC as simplified models of ‘cancer’ and ‘non-cancer’ membranes (DOPC -1,2-dioleoyl-sn-glycero-3-phosphocholine, DOPS – 1,2-dioleoyl-sn-glycero-3-phospo-L-serine). Leakage assays followed the well-established Carboxyfluorescein fluorescence dequenching method as illustrated in **Figure 2a**, and the use of a liquid-handling robot enabled medium-throughput screening of peptides.

**Figure 2.**
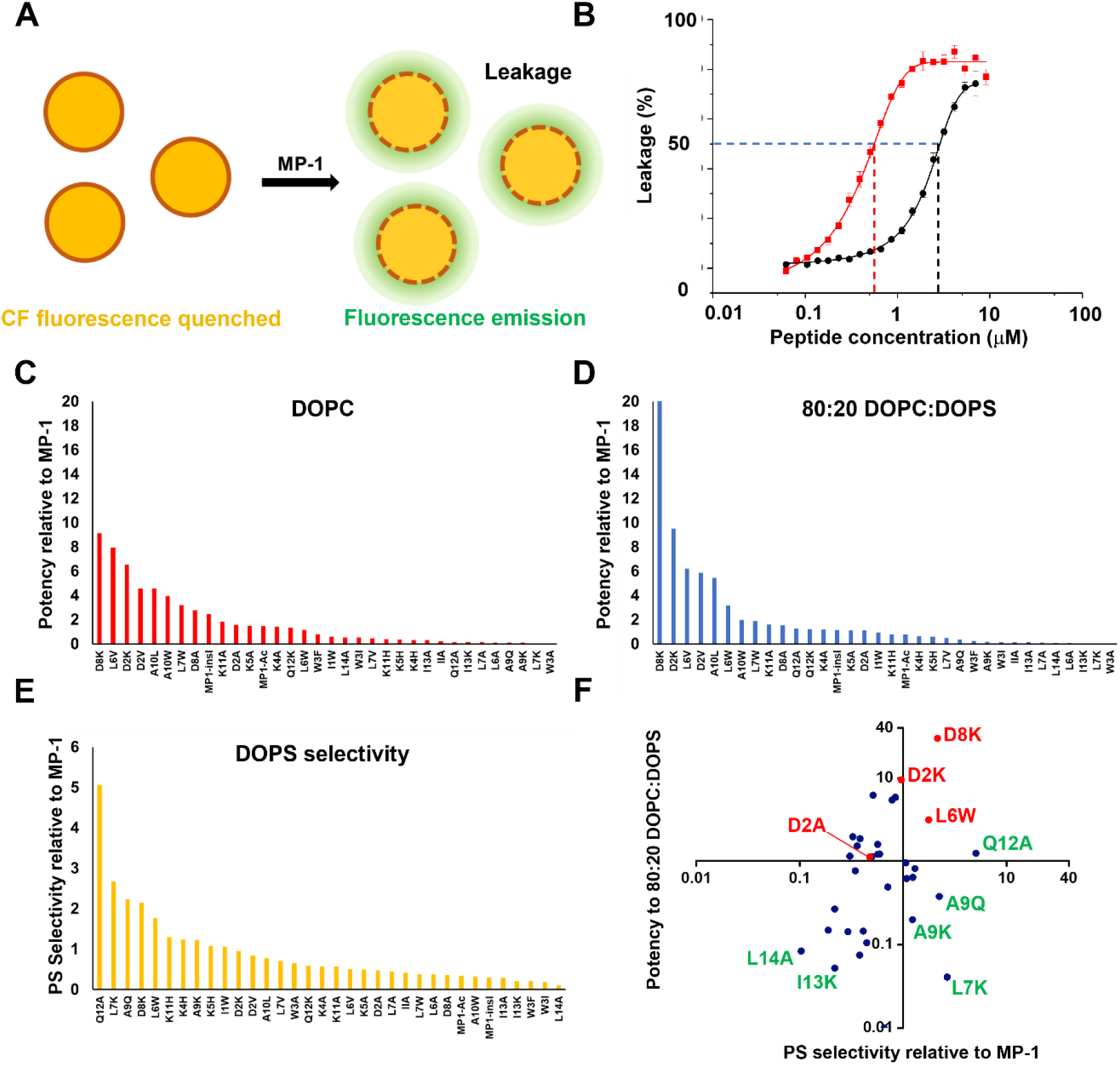
Dose-response analysis of damage to artificial model membranes induced by MP1 and derivative peptides. **a)** Illustration of the principle of the Carboxyfluorescein (CF) leakage assay. CF is encapsulated in artificial vesicles at a high, self-quenching concentration; peptide-induced leakage of CF causes it to dilute in the surrounding media, leading to an increase in fluorescence emission and, therefore, fluorescence intensity is proportional to the degree of membrane disruption. **b)** Example of dose-response leakage data (Black: Red: 80:20 DOPC:DOPS LUVs treated with MP-1): the peptide potency is defined as the concentration of peptide required to induce 50% of the maximal leakage (*L*_50_). **c)** Potency of mutant peptides vs 80:20 DOPC:DOPS LUVs relative to MP-1. **d)** Potency of mutant peptides vs DOPC LUVs relative to MP-1 internal controls, where: 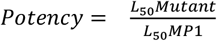. **e)** Selectivity for 80:20 PC:PS membranes vs. DOPC membranes for MP-1 mutants, where: 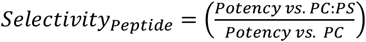. **f)** Plot of peptide potency vs. 80:20 DOPC:DOPS vesicles against peptide selectivity: candidates for further study are labelled in red.

To aid comparison with cell viability data, peptide-induced leakage was quantified in a manner analogous to calculating IC_50_ values; the concentration of peptide needed to induce a fluorescence value 50% of that obtained in the corresponding positive control experiment (vesicles treated with Triton X-100 detergent) this was termed the ‘50% leakage concentration’, see **Figure 2b**. Peptides were assessed against the ‘cancer’ model 80:20 DOPC:DOPS membrane and against the ‘non-cancer’ model DOPC membrane. These data are presented as 50% leakage concentrations in **Table S6**, and also as potency relative to MP-1 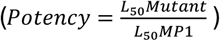 in **Figures 2c** and **2d** allowing easier visualisation of the impact of the mutations. Potency in these two models were also combined to give a measure of relative selectivity for the ‘cancer’ model 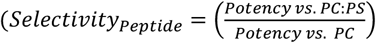, **Figure 2e**. These metrics are convenient to assess the key changes in activity resulting from each specific mutation, relative to the MP-1 peptide. While potency against both of the model membrane compositions are correlated (Pearson correlation value: 0.78464, p = <0.0001), notable ‘selective’ outliers are observed, as shown in **Figure S4**, demonstrating that, for these membrane compositions, it is possible to enhance selectivity to PS without sacrificing ‘on-target’ activity.

Some general observations are readily apparent in **Figures 2c-2e**, and most clearly in which substitutions cause severe loss of potency. Notably, substitutions that reduce hydrophobicity in the terminal residues (I1, I13, L14), and reduction in hydrophobicity in general, reduce potency against model membranes. This reduction is particularly apparent when a K residue is introduced at these hydrophobic positions. Conversely, replacement of negatively charged D residues with positively charged K residues generally increases potency against both membrane compositions. Interestingly, it might be expected that an increase in positive charge of the peptide enhances selectivity to anionic DOPS-containing membranes, but enhanced potency was also observed against zwitterionic PC-only membranes.

A plot of potency against the 80:20 DOPC:DOPS model vs. selectivity, **Figure 2f**, was used to select peptides with improved potency and improved selectivity relative to MP-1 (top right quadrant) as candidates for further study, namely D8K, D2K and L6W. In addition, D2A was selected to take forward into mechanistic studies, due to its comparable potency to MP-1, but reduced selectivity for 80:20 DOPC:DOPS, for possible insights into the mechanism of DOPS selectivity.

### Physicochemical determinants of MP-1 peptide potency and selectivity against model membrane vesicles

Our leakage assay results indicate that bulk physicochemical properties such as charge and hydrophobicity are often not straightforward predictors of peptide activity for MP-1. Similar conclusions have been drawn in previous studies of other membrane-active peptides^16^, suggesting that, in general, membrane-disrupting peptides have a complex structure-activity relationship. Nonetheless, we have attempted to find correlations in our data based on categorisation of the physicochemical changes associated with each mutant as shown in **Figure 3A**. Being composed of only 14 residues, single mutations result in substantial changes to the overall properties of the peptide. Each plot in **Figure 3A** is analogous to **Figure 2f**, with axes corresponding to selectivity and potency. Peptides are separated into categories based on qualitative changes to their physicochemical properties, (e.g. the peptide D8A is categorised as a reduction in negative charge, reduction in steric bulk and increase in hydrophobicity). The ‘positive charge’ and ‘negative charge’ categories are assessed independently in these analyses, to reflect the structural differences implied by addition and removal of K and D residues beyond the net physiochemical charge.

**Figure 3:**
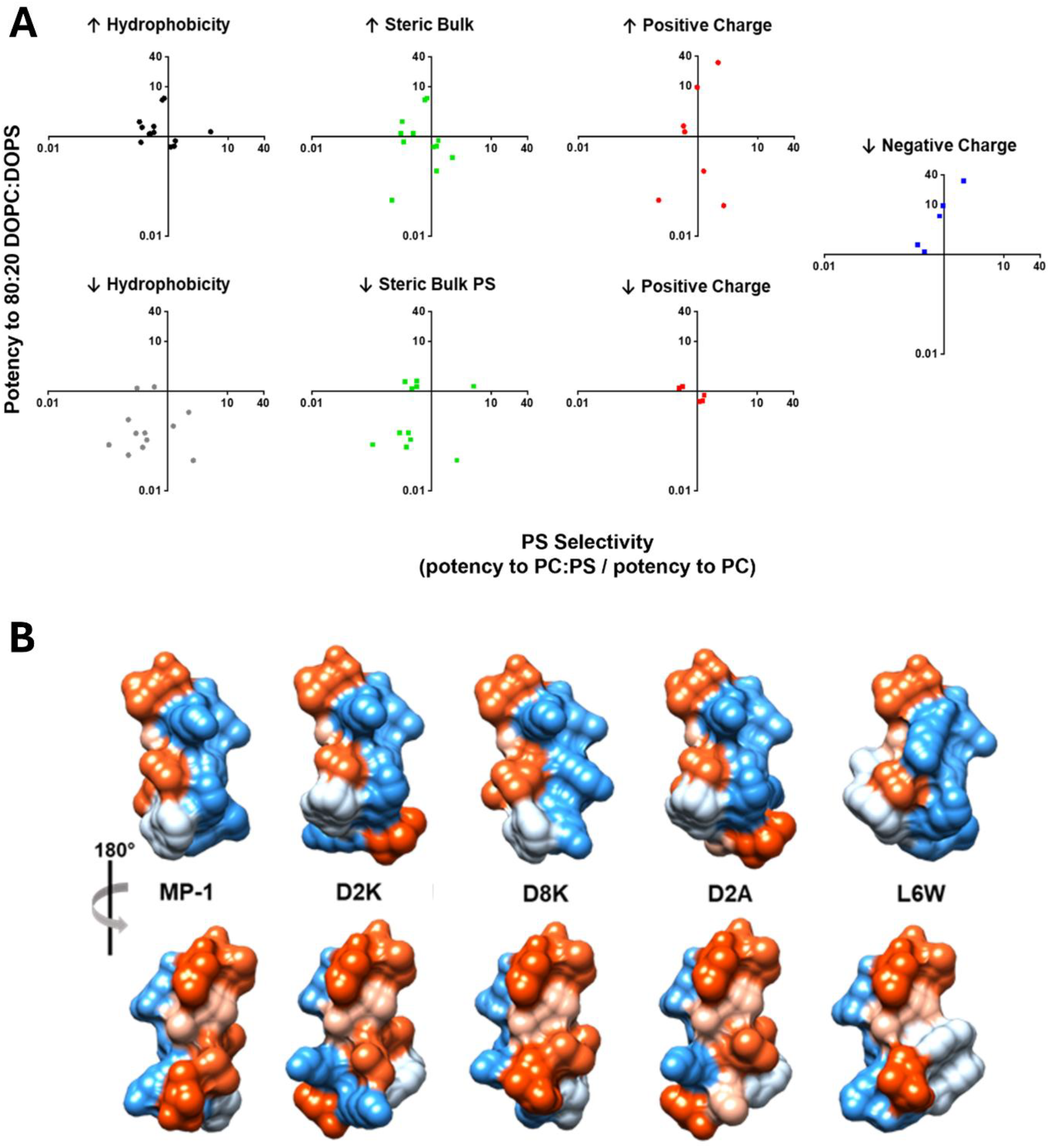
Analysis of physicochemical characteristics of MP1-derived peptides that drive the potency and specificity against model membranes. Structural analysis of peptide mutants. **A:** Plots of potency against 80:20 DOPC:DOPS 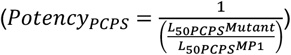 vs. selectivity 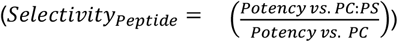 for mutants characterised by a given category of physiochemical change (increase: ↑, decrease: ↓) relative to MP-1. **B**: Kyte-Doolittle hydrophobicity surface space filling models of selected peptide mutants red-hydrophobic, blue-hydrophilic, two projections are shown for each molecule, corresponding to a 180 degree rotation around the axis of the amide backbone, based on *de novo* structure prediction using the PEP-FOLD3 software^17^, surface visualised using USCF Chimera^18^.

As seen in **Figure 3A**, increasing hydrophobicity generally results in an increase in potency, and decreased hydrophobicity results in a decrease in potency, however in both instances, most mutants exhibit reduced selectivity for PS membranes.

The greatest increases in potency against PS membranes were seen for mutants with reduced negative charge (replacement of D residues), but specifically those where D was replaced with K, resulting in an overall increase in charge of +2, likely resulting in increased electrostatically driven adsorption of the peptide to the vesicle membrane in PS containing vesicles. However, of this category, improved selectivity for 80:20 PC:PS over DOPC was only observed in the case of D8K. D2K, while more potent, has comparable PS selectivity to MP-1. Furthermore, reduced potency was observed for three peptides in the increased positive charge category (I13K, A9K and L7K), with I13K also exhibiting reduced PS selectivity. Particularly in the case of A9K, where the reduction in hydrophobicity is less significant, this suggests that in determining PS-selectivity, hydrophobic effects dominate the effect of increased positive charge. Beyond conferring a negative charge, inclusion of PS lipids in a membrane will alter lipid-packing, peptide insertion and packing in the membrane. Many of these mutations also alter the steric properties of the peptide, both locally and in terms of molecular conformation, with clear implications for insertion and packing.

Clearly, changes to bulk physiochemical properties do not provide a full account of the changes in peptide properties resulting from these substitutions. However, categorising mutants according to whether they have an increased or reduced steric volume relative to MP-1, based on R-group size, also does not result in any clear trend in potency or selectivity, leading us to consider the structural effect of substitutions in each of our candidate mutants on the conformation of the peptide molecule as a whole. Intuitively, such changes at a whole-molecule level could have an effect on insertion/packing that would otherwise seem disproportionate to the steric difference at one substitution site. To interpret these changes in behaviour, we explored a 3D molecular modelling approach as shown in **Figure 3B**.

### 3D peptide structure prediction suggests global conformational changes induced by point mutations play an important role in peptide potency and selectivity

We used the PEPFOLD-3 (https://bioserv.rpbs.univ-paris-diderot.fr/services/PEP-FOLD3/) structure prediction software to generate predicted conformations, providing visualisations of possible conformational changes resulting from given substitutions. The PEP-FOLD3 output was used to generate the hydrophobicity surface space-filling models depicted in **Figure 3B**. The PEPFOLD-3 software predicted an α-helical structure for all MP1 derivative peptides. MP-1 is known to undergo a transition from random coil in aqueous media to an α-helical configuration when bound to lipid membranes, which can be probed by circular dichroism spectroscopy^9^. Therefore the predicted structures are likely to best represent preferred configurations of the peptide when bound to the membrane. Changes in orientation of peptide side chains due to changes in the sterics and interactions between nearby amino acids in the peptide will impact how these peptides interact within a lipid membrane environment. While the PEPFOLD-3 software does not explicitly include the liquid crystalline solvation of the membrane environment within its structure prediction, observed changes in side chain orientations are suggestive of changes in the energy balance within the peptide structure that may affect its membrane activity. This analysis also has the potential to group peptides with similar changes in side chain structures to see if they exhibit similar potency in inducing dye leakage from lipid vesicles, where these groupings will likely differ from those previously considered based upon changes in physicochemical properties such as charge or hydrophobicity.

The space filling models of the predicted peptide structures have a more ‘globular’ form than we intuitively expected considering a ‘linear’, α-helical molecule of 14 amino acids, and the volume of individual R-groups make substantial contributions to overall molecular shape. Often differences in side-chain orientations were predicted at positions remote from the substituted residue, for example, the orientation of the W residue R-groups can be seen to change in D8A, which may impact their mode of interaction with the membrane. MP-1 is considered to possess hydrophobic and hydrophilic faces, corresponding to the two projections of each variant depicted in **Figure 3B**. Multiple single point substitutions lead to changes to the overall form of these faces. In the cases of D2K and D2A, the hydrophobic face is predicted to become interrupted by a projecting K residue, with the I1 side chain becoming separated from the hydrophobic patch. Both of these peptides exhibited reduced selectivity for PS membranes, but more notably in D2A where a negative charge is also removed. D8K preserves a MP-1-like hydrophobic face and has improved potency and selectivity, hinting at a role for the hydrophobic/hydrophilic face structure in PS selectivity. This may indicate that the conformation of the hydrophobic face in MP-1, and its implications for insertion and packing, are critical determinants of its PS-membrane selectivity.

The role of the W residue is also notable for both potency and selectivity, as shown in **Figure 2**. Substitutions at W3 generally have a detrimental effect on potency and selectivity, even when replaced with a hydrophobic (I) or aromatic (F) residue. I1W has remarkably similar properties to MP-1, while L6W has improved potency and selectivity. As seen in **Figure 3B**, L6W is predicted to have a conformation with W residues in a ‘stacked’ arrangement, while in I1W, the W aromatic rings have a near co-planar arrangement with W1 protruding onto the hydrophilic face (see **Figure S5**). As a bulky aromatic residue, it is possible that W residues play a role in altering the local lipid packing in the membrane upon insertion, possibly contributing to local induced curvature necessary to initiate poration.

### Mechanistic analysis by AFM reveals multiple modes of action in MP1-like peptides

Taken together, our analysis of membrane permeabilisation data and *in silico* structural analysis of single substitution MP-1 mutants imply a finely balanced, complex relationship between peptide structure, lipid composition and their effect on membrane structural perturbation. AFM studies enable visualisation of supported lipid bilayers (SLBs), with nanometre resolution, over time. Due to the relatively low throughput of AFM studies, a subset of 4 peptides plus MP-1 was taken forwards. D2K, D8K, D2A and L6W were selected for the improvements in selectivity and/or potency relative to MP-1 seen in the analysis of vesicle leakage data and, in the case of D2A, due to its selectivity for non-PS membranes.

SLBs of DOPC and 80:20 DOPC:DOPS were prepared and continuously imaged using a FastScan AFM system enabling an effective frame rate of approximately 2 frames/min for a 1.0 x 1.0 µm^2^ sample area. Representative data for SLBs exposed to each of our selected peptides and MP-1 is presented in **Figure 4**.

**Figure 4.**
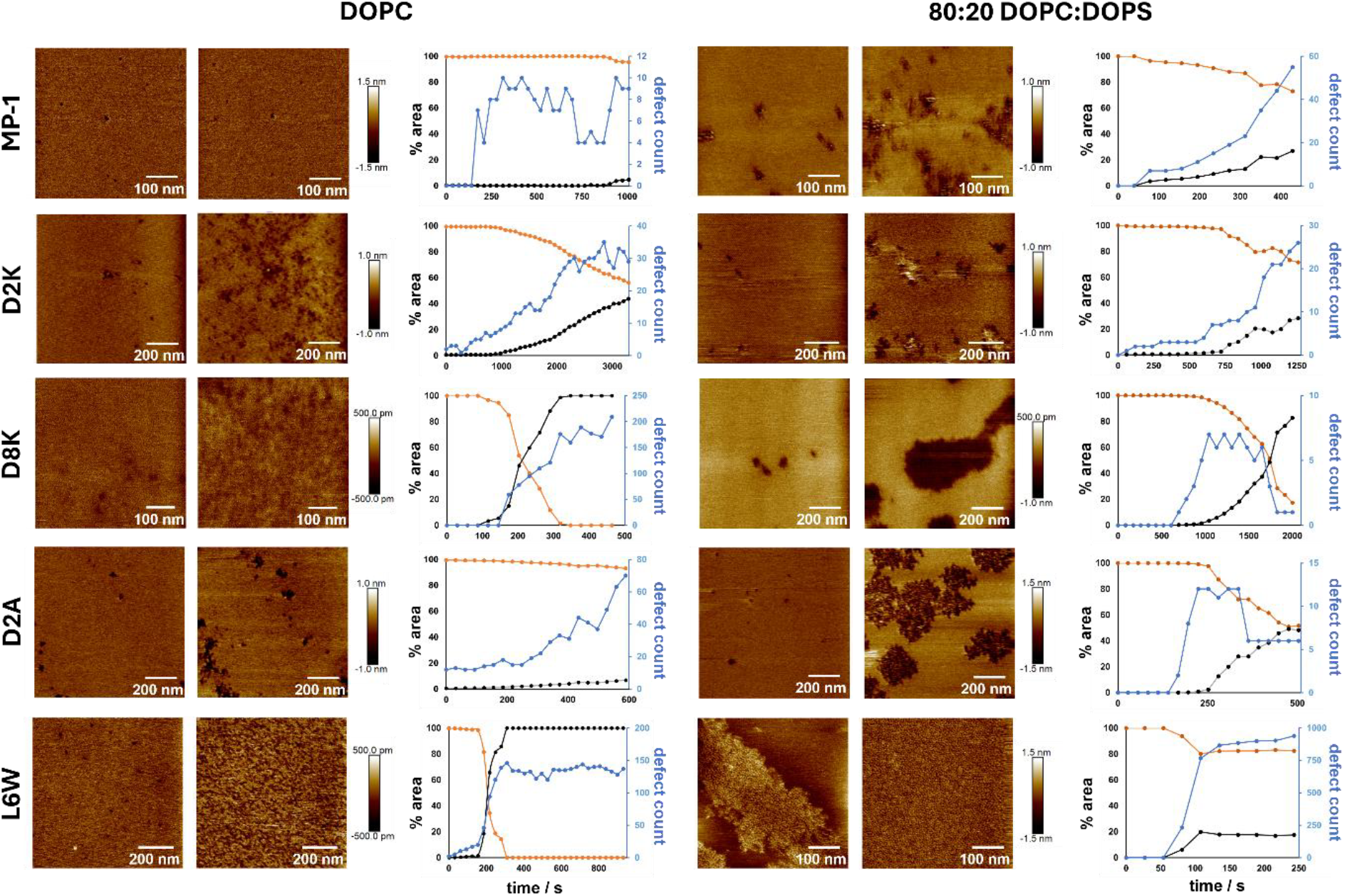
Nanoscale mechanisms of model membrane disruption by MP1-derived peptides. AFM images of DOPC and 80:20 DOPC:DOPS SLBs incubated with 1 γM peptide. Two images are displayed for each combination of membrane and peptide at different time points to illustrate the progression of peptide-induced membrane disruption. The corresponding plots record the % area of the observed region that is intact L_d_ phase membrane 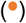 or disrupted phase 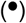, the defect count 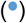 is recorded on the right-hand axis. t=0 is set to the time of peptide addition. Note that it has not been possible to harmonise all axes across plots.

Peptide solutions were added directly to the supernatant solution to give a final concentration of 1 µM initially, although subsequent additions were occasionally made if no membrane perturbation was observed after approximately 10 min. Imaging continued until the membrane perturbations appeared to reach a stable ‘end state’ or, in some cases, until fouling of the probe tip by non-adherent membrane/peptide material made further imaging impossible. **Figure 4** presents two representative images, from a single run, for each combination of peptide and membrane, with time points selected to best illustrate the progression of the particular mode of membrane disruption observed, which differed between peptides studied. Corresponding image analysis data for each run is presented with percentage area of intact membrane and disrupted membrane and the number of distinct ‘defects’.

### PS-containing membranes develop larger defects

The most general observation in these studies was the formation of larger defects in PS-containing membranes. This is observed quite consistently, with the exception of peptide L6W, which exhibits very distinct membrane disruption behaviour (discussed below). The generality of this observation suggests that one of the drivers of PS selectivity seen in MP-1 may, in part, be more severe membrane disruption. This is possibly due to enhanced electrostatic binding of the peptides to the membrane, where 20 mol% PS lipids have previously been shown to enhance membrane partitioning of MP-1 peptides by a factor of 7-8 compared to comparable zwitterionic lipid membranes^9^.

### Distinct mechanisms of action are characterised by the morphology and kinetics of pore nucleation and growth

In **Figure 5** we present diagrammatical representations for the common mechanisms of membrane disruption. DOPC membranes treated with MP-1 exhibit a unique poration mechanism not observed for other peptide-membrane combinations in this study. Discrete uniform pores of a highly regular diameter of approximately 13 nm are observed to form, with many appearing to be transient, persisting for approximately 2 frames of imaging time (∼40 s), although a smaller number of more stable pores were also observed. The pores observed in this system did not undergo any growth or morphological change. We classify this mechanism as ‘small transient pores’ (**Figure 5 A**). Contrasting this with MP-1 vs. 80:20 PC:PS membranes, we see a ‘progressive poration’ (**Figure 5 B**) mechanism, where pores of varied diameter and morphology nucleate and undergo some limited growth, with continuing nucleation eventually leading to a membrane with a high percentage of the observed area being composed of ‘disrupted phase’, that being the membrane type which comprises these defects.

**Figure 5:**
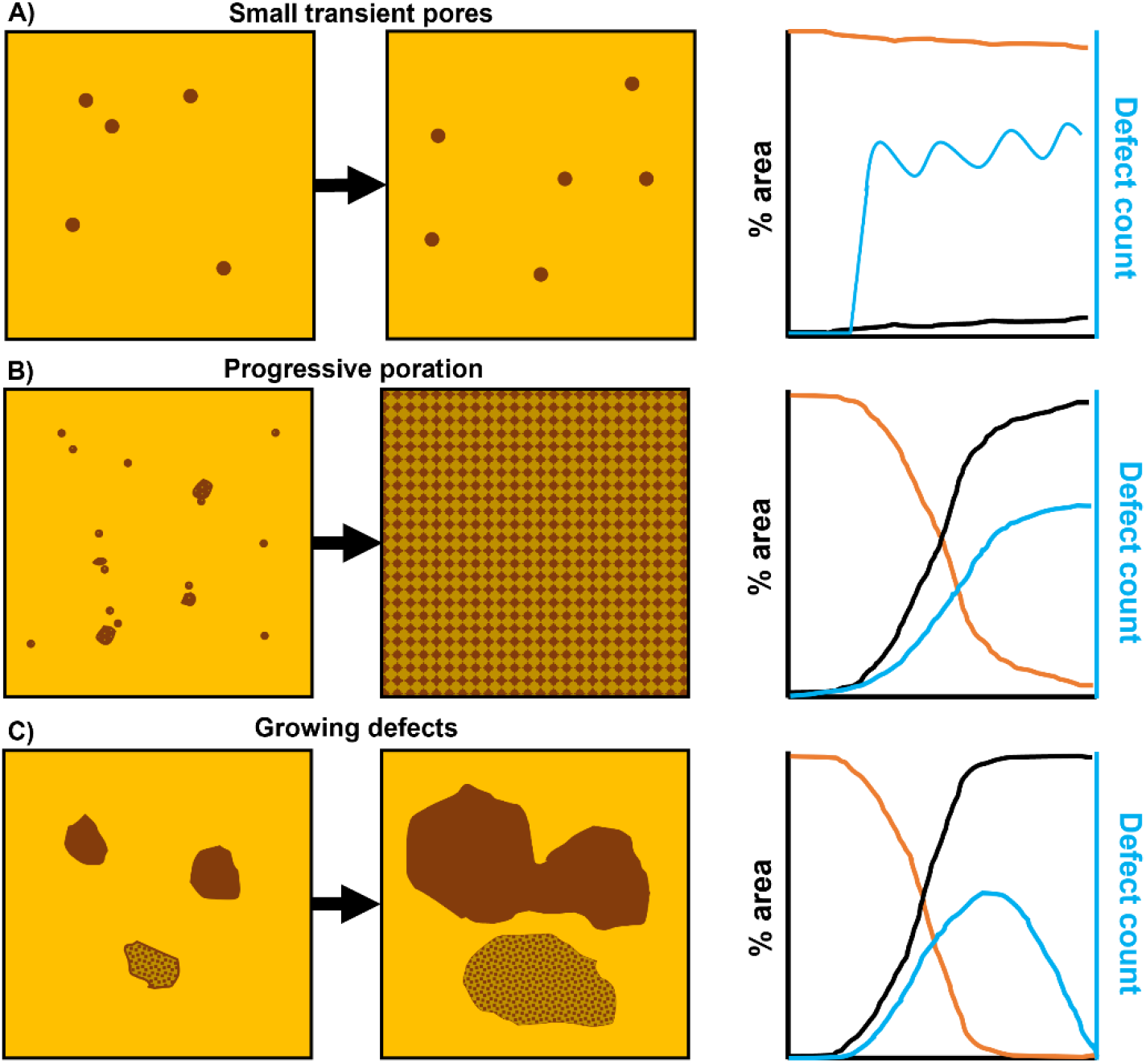
Diagrammatical representations of mechanisms of peptide action on SLBs. **A)** Small transient pores of uniform size (≈13 nm). **B)** Progressive formation of small stable pores leading to widely disrupted surface. **C)** Widespread disruption resulting from the growth and merger of large defects

This second mechanism of ‘progressive poration’ is the one most commonly observed. For example, D2K follows this mechanism for both membrane compositions, with larger defects forming in the PS-containing membrane. This mechanism is also applied to D8K, D2A, L6W vs. DOPC membranes. The third general mechanism was observed with D8K and D2A. ‘Growing defects’ (**Figure 5 C**) are characterised by fewer pore nucleation events with rapid growth, particularly in the case of D8K. This mechanism is most likely to reach an end point when the membrane in the imaging area is fully disrupted. The defect count in this scenario thus tends to reduce over time as previously separate defects merge. Our illustration in **Figure 5 C** also depicts two different types of defect area / disrupted phase. Some disrupted phases possess a relatively smooth, uniform surface, such as in D8K against 80:20 DOPC:DOPS, while others, such as D2K possess height irregularities. Imaging of the latter type was often subject to fouling of the AFM probe tip, suggesting that lipid or lipid-peptide aggregate material is lost from the surface.

The action of L6W against DOPC membranes follows the most commonly observed progressive poration (**Figure 5 B**) mechanism. However, in some instances a dynamic filamentous structure was observed. Although the structure cannot be definitively identified from these images, its appearance informs our interpretation of anomalous behaviour of L6W against PS-containing membranes. In these experiments, no membrane disruption or poration was observed over a long period of imaging. Due to this lack of activity, a subsequent aliquot of L6W was added to the sample to give a peptide concentration of 10 μM. After several minutes, the appearance of plaque-like height anomalies was observed. These features evolve rapidly into an ordered network which rapidly fills the observed area. This structure can somewhat resemble the end state of the progressive poration mechanism due to the dense pattern of roughly circular depressions with a regular diameter of ∼6.1 nm and a depth of ∼7 nm, while the width of the ‘filaments’ / rims of the depressions is approximately 3 nm. Taken together with the observed filamentous structures induced by L6W on DOPC membranes, we interpret this structure as a network of tubules or other linear, filamentous structures rather than an expanding area of dense poration.

We assume that these structures are a network of self-assembled peptide or possibly locally-curved membrane, induction of membrane curvature being a key part of the action of MP-1 and other membrane disrupting peptides.

The addition of a second Tryptophan residue in L6W, is predicted in our molecular modelling (**Figure 3B**) to result in a dominant conformation where the aromatic moieties of the W6 and W3 Tryptophan side-chains are arranged in a stacked conformation that implies a π-stacking interaction. Such interactions, along with Π-cation interactions, are known to play a role in intermolecular interactions of peptides^19,20^, and W residues have been shown to be of critical functional importance for a number of membrane-disrupting peptides^21,22^.

### Relating mechanism to leakage assay data

There are a few general observations in our AFM studies, for which it is tempting to infer a relationship between a mechanistic change and the performance of these peptides in our leakage assays.

The general observation of larger defects in PS-containing membranes is consistent with the increased potency of MP-1 (and most of its mutant derivatives) in leakage assays against this membrane composition.

In the DOPC experiments, the highly potent mutants D8K and D2K are notable for the very high density of defects they are able to induce compared to MP-1. A similar end-state is observed in DOPC vs. L6W, although this must be interpreted more cautiously. This contrasts with D2A vs. DOPC, where nucleation of defects progressed much more slowly, and, uniquely for peptides against DOPC membranes, appears to possess a mechanistic character combining ‘progressive poration’ and ‘growing defects’ (**Figure 5 B/C**), with continuous nucleation of new defects at a slow rate, but those defects show significant growth over time. Its end state more closely resembles those observed for MP-1 and D8K against PS-containing membranes. However, D2A is unique in the peptides taken into AFM studies in that it has a reduced PS-selectivity relative to MP-1 (due to enhanced activity against DOPC). This may therefore reflect the convergence in membrane disruption mechanism we observe in our AFM studies and the ‘growing defects’ mechanism observed against SLBs may represent a more disruptive mechanism in a vesicle membrane.

D8K, which is the most potent peptide in our leakage studies, exhibits exceptional potency against PS-containing membranes, where it causes rapid growth of defects. This mechanism progresses to an end state where the majority of the observed membrane is of the ‘disrupted phase’. The activity of D2K and D2A against PS or non-PS membranes is more similar and their mechanisms observed by AFM are also consistent. Contrastingly, D2K’s action against PS-containing membranes is more comparably to MP-1 (growing defects) but pores were significantly smaller and slower growing than those of D8K. The behaviour of D8K is also distinguished from that of others against PS-membranes in that the defects are free of the height irregularities discussed earlier, which correlated with tip fouling, and we therefore tentatively identify as lipid/peptide material protruding or detaching from the membrane surface. How this might relate to enhanced potency in vesicle studies is unclear but further indicates a divergence in mechanism.

The anomalous activity of L6W was not anticipated based on leakage assay data, it was selected for further study purely based on its enhanced potency and PS-selectivity relative to MP-1. That L6W performs well against DOPC membranes is not surprising given the rapid ‘progressive poration’ mechanism. The structures observed for L6W in out PS-membrane experiments is harder to interpret as it is not clear if they contain defects comparable to those in our other experiments. The high potency of L6W in our leakage assays might be used to inform our interpretation of these structures as representative of a membrane perturbation sufficient to induce leakage in an unsupported vesicle membrane. However, it is possible that the behaviour is different in SLB systems, as it is conceivable that the solid substrate imposes a constraint on the development of L6W-induced membrane perturbations.

### Cytotoxicity of MP1-derivatives mostly correlates with model membrane studies but with nuance that indicates broader cell-specific properties are also important

In **Figure 1**, we selected a panel of cell lines to represent relatively ‘resistant’ cells (BT-474, MCF-10A) and relatively ‘sensitive’ cells (MDA-MB-468, AU565). One goal of selecting cell lines with different extremes of MP-1 responsiveness was to assess whether these differential responses would be maintained with key mutant peptides, and whether cells would reproduce the trends seen with synthetic membranes (Figures 2 and 4). Selected mutants identified in leakage assay screening (**Figure 2**) were tested against these cell lines to assess their activity in the more complex cellular context. IC_50_ values for D2K, D8K, D2A and L6W were determined in cell viability assays in all four cells lines, and are presented alongside the values for MP-1 in **Figure 6A**.

**Figure 6:**
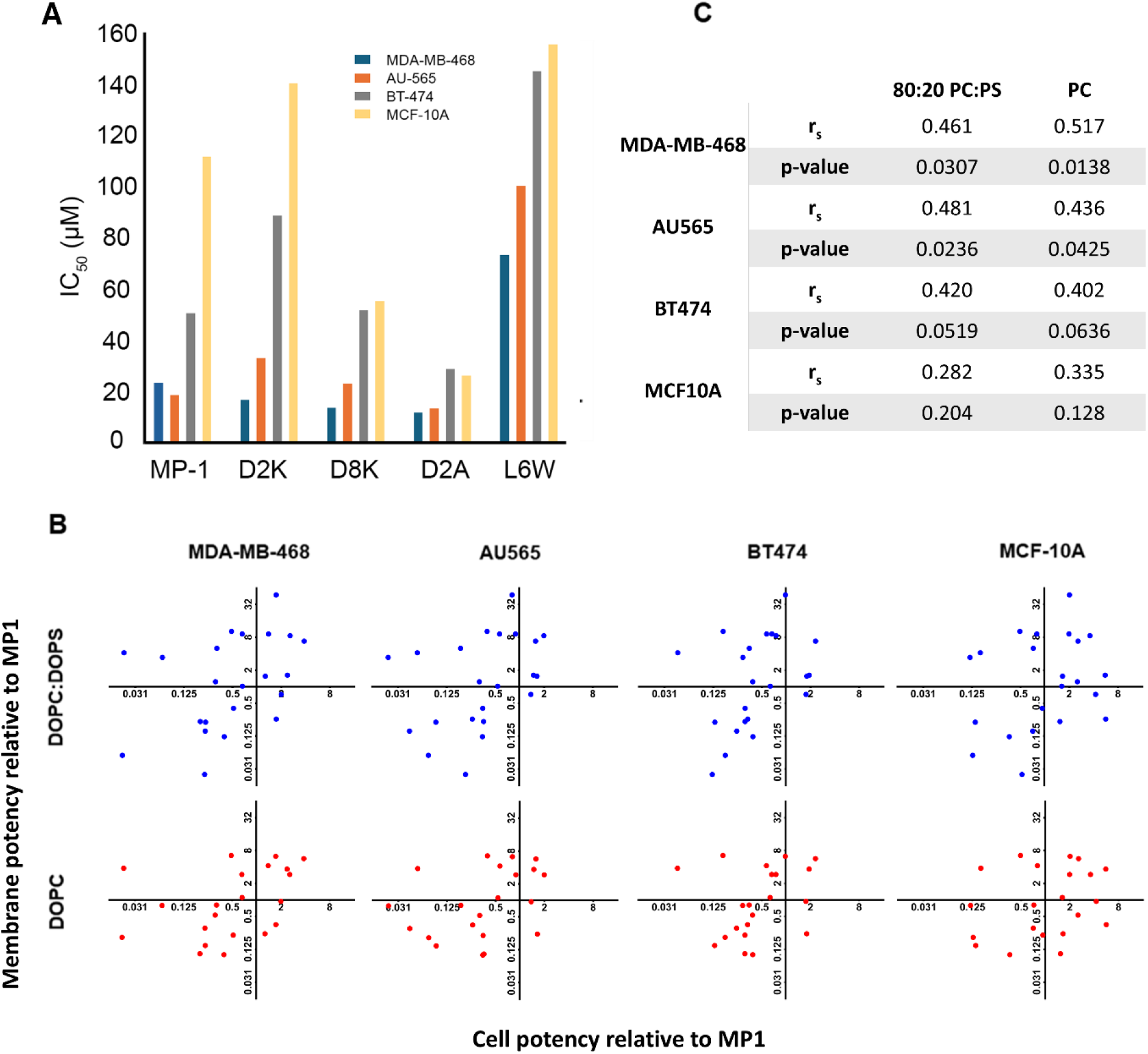
Correlation between cell viability and model membrane leakage assays. **A**: IC_50_ values for selected peptide mutants and cell lines. **B**: Scatter plots of potency values 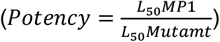. for all studied peptide mutants versus each model membrane (y-axes, **DOPC**: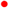, **80:20 DOPC:DOPS**: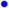) and against the different cell lines (IC_50_(MP1)/IC_50_(mutant)) (x-axis), Log_2_ scale. **C**: Spearman analysis of data in **5B**. r_s_ – Spearman correlation coefficient.

D2A has greatly reduced IC_50_ values for the ‘resistant’ cell lines (blue and grey bars in Figure 6A) and much smaller reductions in IC_50_ values in ‘sensitive’ cell lines (blue and red bars). In our prior leakage assays (**Figure 2**), D2A was selected for further study because of its reduced PS-selectivity, driven by enhanced potency against DOPC membranes and negligible improvement in potency against DOPC:DOPS 80:20. This appears to be reflected directly in the increased potency (reduction in IC_50_ values) in resistant cell lines seen, with similarly negligible improvement in the sensitive lines (**Figure 6A**), although this differential in cell lines is even more striking. The AFM mechanistic studies detailed in **Figure 4**, did indicate a divergence in mechanistic behaviour for D2A where behaviour against DOPC membranes more closely resembled activity against PS-membranes by other peptides.

The observed IC_50_ values for D2K and D8K are correlated with what we might anticipate based on our leakage data, an overall reduction or similar IC_50_ values to MP1 for most cell lines. D8K is the most potent peptide against MDA-MB-468 cell lines but its increase in potency against the MP1-resistant MCF-10A cell lines is more significant, making this peptide less selective than MP1. This reduction in IC_50_ for ‘resistant’ cell lines is somewhat greater than we might have expected based on leakage data, however due to the complexity of cell models, both in terms of membrane composition and other non-membrane factors influencing survivability, a very close correlation should not be expected. Importantly, however, for D2K we observe both a decease in IC_50_ against the sensitive, cancer cell line MDA-MB-468 and increase against the resistant, non-cancer cell line MCF10A. Overall, this makes D2K more selective than MP1 for these two cell lines, which makes it an interesting candidate peptide to progress alongside MP1 in future work. It should be noted that this increased selectivity is nuanced, as D2K is less potent (and therefore less selective) than MP1 against the other two cancer cell models: AU-565 and BT474. Therefore, as could be expected, this improved potency and selectivity is dependent on phenotypic properties of specific cancer cell lines.

The behaviour of L6W is perhaps the most poorly aligned with our leakage assay results, being a potent and PS-selective peptide in those experiments and displaying an unusual combination of changes to IC_50_ values, being less potent against both ‘sensitive’ cell lines and BT-474 cells, while being more potent against MCF-10A cells. AFM studies with L6W indicate that it has a highly divergent mechanism contrasting with all other peptides studied, therefore we might conclude that this mechanism of activity is less effective in cell membranes. Furthermore, the AFM results may indicate peptide self-assembly in L6W, it also is possible that, in the cell culture environment, such behaviour is a greater hindrance to membrane activity.

With the partial exception of L6W, the overall potency of the selected peptides is consistently reflected in the model membrane leakage and cell viability assays. Further refinement of the membrane models, with better compositional and structural alignment to cell plasma membranes should enable us to extract more detail relating peptide activity and membrane composition, to better understand the membrane properties that are driving selectivity in sensitive and resistant cells. The potential complicating factor of other cellular structures and processes, such as membrane repair activity, might be investigated with a bioinformatics approach to further enrich the findings.

### Closer biomimetic membrane models are required to strengthen the correlation between model membrane disruption and cell-line cytotoxicity

In **Figure 6 B**, scatter plots were generated for all peptide mutants for which cell viability assays were performed. The relative potency of peptide mutants in model membranes is compared with their relative potency against each cell line to look for those combinations of model membrane and cell lines where the activity of the peptides is best correlated. A Spearman’s correlation test is presented in **Figure 6 C**. The r_s_ (Spearman correlation coefficient) values obtained generally indicate moderate correlations for the two sensitive cell lines (MDA-MB-468 and AU-565) and also the BT-474 cell line, which is intermediate between the sensitive cells and the highly resistant MCF-10A cells. In comparison, there is only a weak correlation for the resistant MCF-10A cells. Furthermore, the correlations for the two sensitive cell lines are statistically significant (p<0.05), with the intermediate BT-474 cells being close to this threshold of significance, but the correlation for the highly resistant MCF-10A cells is not found to be significant. Taken together, this indicates that our model membranes are more reliable for determining the effect of these peptides against cell lines sensitive to MP1 but do not capture the resistance mechanisms of cells that exhibit low sensitivity to the MP1 peptide. Therefore the model membrane screening approach is more attuned to identifying novel peptides of increased potency, but less reliable for predicting peptides that might have improved selectivity for sensitive over resistant cells.

The stronger correlation between cell and leakage studies for sensitive cell lines may indicate a better correlation in the membrane composition or physical properties between those categories, where the model membranes in this study contain unsaturated lipids in the L_α_ phase, which are loosely-packed in the membrane compared to vesicles containing saturated phospholipids and/or cholesterol. Indeed, tightly packed regions of membrane may be more important for modelling the mechanisms of cell resistance to MP-1 and its derivatives, where much poorer correlations are observed for resistant cell lines. However, other native membrane structural features such as membrane asymmetry or electrochemical membrane potentials may also play an important role. Furthermore, active cellular protection/repair mechanisms are likely less prevalent in sensitive cell lines, making their behaviour easier to model with passive model membranes. Model membranes with improved composition or physical properties, more similar to native membranes, could provide more accurate screening, within the limitations of a model lacking cellular processes such as membrane repair or structures such as the glycocalyx. However, this identifies an opportunity to add further complementary biological assays downstream of model membrane screening and add value to the complete data set. Overall, these analyses support the conclusion that the model membrane screening approach is a valid methodology for identifying candidate peptides, particularly for improved potency against sensitive cell lines, but requires refinement in terms of model membrane composition and to control for other cellular properties, which may have a stronger influence in the outcomes for cells that have a high resistance to the activity of MP1.

## Conclusions

Development of cancer therapeutics derived from membrane disrupting peptides is currently held back by insufficient understanding of their mechanism of action. We have therefore developed a hybrid approach to rapidly screen candidate peptides based on MP-1. We find substantial variation in the membrane activity of MP-1 mutants, allowing us to take a selection forward to biological assays of activity and biophysical studies to develop our understanding of the mechanism of membrane disruption. These findings can then be fed back to refine our assays through a design-test-learn cycle, e.g. through better biomimetic model membrane compositions.

MP-1 is capable of disrupting cellular membranes leading to cell lysis, with the extent of activity varying significantly across the range of cell lines studied. There was not a simple selectivity for cancer-derived cell lines, but sensitive and resistant categories were identified as a basis for our combined biological and biophysical screening approach. Differences in membrane permeabilization between these categories suggests a different mechanism of MP-1 activity or cellular response.

Rapid screening of mutants in our LUV leakage assays and subsequent cell viability assays indicate that the membrane disrupting activity of MP-1 is finely balanced, with single residue mutations able to greatly enhance or disrupt its overall potency and selectivity for particular model membrane compositions or cell types. Notable findings include the essential nature of the I1, I13 and L14 terminal hydrophobic residues and D to K replacement boosting activity against both membrane compositions. No strong indication as to the structural basis of PS-selectivity was apparent in these results. This led us to investigate the role of molecular conformation by computational modelling. Using the PEPFOLD-3 structure prediction software, we observed how single point mutations could radically alter the predicted conformation of the molecule and the amino acid side chains in particular, at positions remote from the mutated site due to changes in the global energy balance and interactions within the peptide. Mutants that preserved a MP-1-like geometry on their hydrophobic face were found to be more potent (compare D2K and D8K). We theorise that these differences in the most favourable conformations drive membrane selectivity through preferential insertion and packing in membranes of particular lipid compositions.

The exact mechanism of disruption in model membranes was also found to vary considerably with these single point mutations, as observed directly by AFM. Transient and highly uniform pores were observed in DOPC SLBs exposed to MP-1, while D8K induced rapidly expanding regions of disruption in membranes of the same composition. Generating more, or larger size, defects correlates well with potency observed in leakage assays. This suggests mechanistic differences can drive changes in activity. Leakage assay data on our selected mutants were found to correlate with corresponding cell viability assays, validating the methodology. This correlation is moderate and statistically significant for cell lines that are sensitive to the action of MP1, but only moderate-to-weak for resistant cell lines with no or weak evidence to reject the null hypothesis of there being no correlation. This implies our current model membrane screening approach is valid for identifying peptides with improved potency, but is unable to reliably predict whether a peptide may have improved selectivity for sensitive cells compared to resistant cell lines, as these assays are poor predictors for the peptide’s effect on the latter. This reinforces the need for refinement of the model membrane systems and supplementation with other methods to investigate the interaction of these peptides with other cellular components and processes.

We have found that MP-1 derivatives are able to induce disruption in model membranes and cell lysis in a manner that can be highly selective for particular cell types or membrane compositions and is highly sensitive to single mutations in its amino acid sequence. Such mutations are capable of radically altering potency, selectivity and mechanism of membrane disruption, but the effect of a given mutation cannot be easily predicted, with changes to molecular conformation likely playing a role.

The findings presented here provide a basis for further refinement of our approach and indicate that rational mutation of MP-1 to enhance characteristics desirable in a targeted cancer therapeutic is achievable. For example, the D2K mutant peptide showed enhanced potency and selectivity compared to MP1 when comparing activity against the sensitive MDA-MB-468 (a triple negative breast cancer cell line derived from a metastatic tumour) and the resistant MCF10A (a non-malignant breast epithelial cell line), and is therefore a good candidate for further investigation alongside MP1.

## Materials and methods

### Materials

Peptides were synthesised by Bio-Synthesis Inc., Lewisville, TX, USA to >95% purity with counterion exchange to HCl, Peptides were lyophilised and shipped on dry ice. Quality control was conducted by HPLC and mass spectrometry by Bio-Synthesis Inc., and we checked selected peptides for purity by analytical HPLC (see SI **Figure S8**). All lipids; DOPC (1,2-dioleoyl-sn-glycero-3-phosphocholine) and DOPS (1,2-dioleoyl-sn-glycero-3-phospho-L-serine (sodium salt)) were obtained from Avanti Polar Lipids Inc. (Alabaster, AL, USA). NaCl, HEPES buffer, were all obtained from Sigma-Aldrich Company Ltd. (UK).

## Methods

### Production of large unilamellar vesicles (LUVs)

A lipid solution (1 mL, 15 mM) of the desired composition was dried under a steam of nitrogen, to a thin film, further dried under vacuum and rehydrated in 5(6)-Carboxyfluorescein (CF) solution (120 mM in 10 mM HEPES pH 7.4) followed by 5 freeze-thaw cycles in liquid N_2_. The suspension was then extruded through a 400 nm polycarbonate membrane using an Avestin LF-1 extruders (biopharma process systems, UK) and unencapsulated CF was removed by size exclusion chromatography using PD-10 gel filtration columns (GE Healthcare Bio-Sciences AB,

Uppsala, SE). Final lipid concentration was determined by colourimetric total phosphorus assay^23,24^.

### CF leakage assays

A custom program was written for a Hamilton Microlab Star M liquid handling robot (Hamilton robotics Ltd.) wherein a serial dilution of peptide solution into HEPES (pH 7.4 or 6.5, 10 m) buffer was performed, followed by a fixed concentration of CF-loaded vesicles. Negative and positive controls were established by addition of vesicles to peptide-free buffer and buffer containing 0.6 mM Triton X-100. The final assay plate (384-well black OptiPlate, PerkinElmer LAS (UK) Ltd) was transferred to a Perkin Elmer Envision plate reader where the CF fluorescence intensity (*λ*_ex_: 495 nm / *λ*_em_: 517 nm). Peptide stock solutions were prepared fresh from lyophilised powder dissolved in the desired buffer and concentration was determined by tryptophan absorbance using a Nanodrop spectrophotometer, measuring the characteristic absorbance at 280 nm.

Serial dilution of the peptide was used to achieve solutions with final peptide concentrations of between 0.01 and 1 γM, to which were added a fixed aliquot of vesicles, to a final concentration of 20 γM, plus a control where no peptide is added to the vesicles and a ‘100 % release’ control, where an aliquot of Triton-X100 is added instead of peptide to a final concentration of 0.06 mM. CF fluorescence was quantified after incubation for 5 minutes. Each experiment was carried out in triplicate. Release was calculated as a percentage of the 100 % release value after subtraction of the background fluorescence of the no peptide control. 50 % leakage values were calculated by fitting the log plot of leakage % against peptide concentration using a sigmoidal dose-response fit.

### Molecular modelling

Peptide 3D structures were generated using the PEP-FOLD3 program hosted at https://bioserv.rpbs.univ-paris-diderot.fr/services/PEP-FOLD3/^17^. This program performs *de novo* structure prediction of short chain peptides. This prediction was then used to generate hydrophobicity map space-filling models to assess the geometry of MP-1 and its derivatives in the USCF Chimera software^18^.

### Preparation of Supported Lipid Bilayers (SLBs)

Dried lipid films of the desired composition were rehydrated in MilliQ water to give 1 mL of solution at a lipid concentration of 0.5 mg/mL, suspension was achieved by thorough vortex mixing. The resulting suspension was subjected to 5 minutes of sonication using a probe-type sonicator, or until the solution appeared less turbid. The resulting suspension (75 mL) was applied to freshly cleaved mica substrates and incubated at 40 °C in a petri dish containing a small piece of wet tissue to prevent drying out. After 1 hour, 25 mL of a 20 mM MgCl_2_ solution was added to each substrate and incubated for a further 30 minutes at room temperature. Immediately prior to use, MilliQ water (30 mL) was added to the substrate, followed by 8 washings of the surface by sequential addition and removal of further 30 mL aliquots of MilliQ water using a pipette. The final volume of supernatant solution during imaging was approximately 200 mL.

### Atomic Force Microscopy (AFM) of Supported Lipid Bilayers (SLBs)

AFM experiments were conducted using a Bruker dimension FastScan instrument and FastScan-D probes operating in tapping mode. Typically, a 1 mm x 1 mm area was imaged continuously throughout the experiment. After imaging the membrane for some time to observe that it has a stable morphology, an aliquot of peptide solution of approximately 10 mL volume is added directly to the supernatant solution during imaging, to give a final peptide concentration in the solution of 1 mM. Imaging was continued until the development of peptide-induced changes was complete. Occasionally, imaging was disrupted by lipid material coming away from the surface due to peptide-induced disruption, although this generally only occurred once disruption was well advanced. To maintain good condition, probes were rinsed thoroughly with dilute detergent solution, milliQ water, and finally isopropanol, followed by drying under a stream of nitrogen, failure to do so led to rapid deterioration due to contamination with lipid / peptide material.

### Cell culture

Cells were originally obtained from ATCC and were subjected to mycoplasma testing and STR typing (Source Bioscience, UK) before use. HepG2, PC-3, BT474, AU565, MDA-MB-468, MDA-MB-231, MCF-7, HB-2, HEK-293, WI-38, HFFF2 and 3T3 were all grown in DMEM medium (Thermo Scientific, Waltham, USA) supplemented with 10% fetal calf serum (FCS; Thermo Scientific, Waltham, USA) and 100 units/ml of penicillin/streptomycin (ThermoScientific, Waltham, MA, USA). Whereas MCF10A cells were grown in DMEM/F12 (Thermo Scientific, Waltham, USA), containing 10% horse serum (Thermo Scientific, Waltham, USA), 20 ng/ml epidermal growth factor (PeproTech, UK), 500 µg/ml Hydrocortisone (Sigma-Aldrich, UK), 500 ng/ml Cholera Toxin (Sigma-Aldrich, UK), 10 µg/ml Insulin (Sigma-Aldrich, UK) and 100 units/ml of penicillin/streptomycin (ThermoScientific, Waltham, MA, USA). Cells were cultured and maintained in a humidified incubator with 5% CO_2_ at 37°C. All experiments were conducted at cell densities that allowed exponential growth or mentioned otherwise.

### MTT assays

Bioactivity of peptides was evaluated on cell lines using MTT assays^25^. AU565, MDA-MB-468, BT474 and MCF-10A cells were seeded in 96 well plates with respective growth medium containing 10% FCS at 1×10^4^ cells per well and incubated for 18 h. Cells were then treated with various concentrations (0 – 100 µM) of MP-1 or mutant peptides for 24 h in growth medium without FCS. Cells were then incubated with 0.5 mg/ml MTT (Thermo Scientific, Waltham, USA) dissolved in growth medium without FCS for 3h. Formazan crystals were then dissolved in isopropanol and absorbance was measured at 570 nm using a spectrophotometer microplate reader (Shimadzu, UK).

### Membrane permeability assay

Cells were seeded in 24 well plates at 2.5×10^4^ cells/well for 18h. Then cells were treated with MP-1 peptide at the respective IC_50_ doses for 24h. Cells were stained with 5 µg/ml Hoechst 33342 (Sigma-Aldrich, UK) in phosphate buffer saline (ThermoScientific, Waltham, MA, USA) for 30 min, and further with 1 µg/ml propidium iodide (Sigma-Aldrich, UK) for 5 min before observing under the fluorescent microscope (EVOS FL auto, Thermo fisher Scientific, UK).

### Whole cell patch clamp recording

MCF10A and MDA-MB-468 cells were grown on glass coverslips for 24h prior to whole cell patch clamp recording to measure the plasma membrane potential^26^. Current clamp recordings were made using an Axopatch 200B ampiflier (Molecular Devices) at room temperature. Data were digitized using a Digidata 1440A interface (Molecular Devices), low-pass filtered at 10 kHz and analysed using pCLAMP 10.4 software (Molecular Devices). The standard extracellular physiological saline solution (PSS) contained the following components (in mM): 144 NaCl, 5.4 KCl, 1 MgCl_2_, 2.5 CaCl_2_, 5 HEPES, and 5.6 glucose adjusted to pH 7.2 with NaOH. For the Na^+^-free PSS, NaCl was replaced with 144 mM N-methyl-D-glucamine (NMDG) or choline chloride (ChoCl) adjusted to pH 7.2 with HCl. The standard intracellular (pipette) solution contained (in mM) 5 NaCl, 145 KCl, 2 MgCl_2_, 1 CaCl_2_, 10 HEPES, and 11 EGTA adjusted to pH 7.4 with KOH. In the mode ‘I=0’, a steady V_m_ was recorded within 5 s of achieving the whole cell configuration. Cells were held at the whole cell configuration for up to 10 min with constant flow of fresh PSS and the V_m_ was recorded. MP-1-containing PSS was directly perfused onto the cells using a ValveLink 4-channel gravity perfusion controller (AutoMate Scientific) at a rate of ∼1.5 ml/min. The solution was allowed to equilibrate for ∼4 min following switching prior to measurement at steady state.

Thus, the mean V_m_ over the last 5 s in control vs. MP-1-containing PSS was used to compare the V_m_ data. The V_m_ signals were sampled at 200 Hz. Liquid junction potentials were corrected using the integrated tool in Clampex 10.4.

### Mitochondrial membrane potential

JC-1 is a membrane permeant dye used to monitor the mitochondria membrane potential (MMP). Cells were grown in 24 well plate (2.5 x 10^4^ cells/well) in complete medium for 18 h. MP-1 was treated to the cells at respective IC_50_ dose for 24h. For measuring the MMP, post treatment with MP-1, cells were incubated with 5 µM JC-1 dye (Thermo fisher Scientific, Waltham, MA, USA) for 30min at 37°C. Then cells were analysed under a fluorescence microscope (EVOS FL auto, Thermo fisher Scientific, UK) with an excitation wavelength of 488 nm and an emission range between 515/545 nm and 575/625 nm respectively.

### Apoptosis (annexin V) assays

After treatment of cells with respective IC_75_ dose of MP-1 for 24h, cells were trypsinised and washed with ice cold PBS and suspended in annexin binding buffer (Thermo fisher Scientific, Waltham, MA, USA). Annexin V-FITC (Thermo fisher Scientific, Waltham, MA, USA) at a final concentration of 2 µg/ml was added to the cells and incubated for 15 min under dark conditions. Prior to flow cytometry, 2 µg/ml 7-AAD (Thermo fisher Scientific, Waltham, MA, USA) was added and cells were analysed using a CytoFLEXS flow cytometer (Beckman, California, USA). The data was analysed using the FlowJo software (version 10.6.1).

### 3D spheroid assay

Low adherent round bottom 96 well plates were used to culture the spheroids. 1000 cells/well were added with 180µl of growth medium supplemented with 10% FCS along with 2.5% matrigel matrix (Corning, USA). The 96 well plates containing the cells were then centrifuged for 10mins at 350g and then then incubated at 37°C with 5% CO_2_ for 48h. Post incubation, the tumor spheroids were treated with IC_50_ dose of MP-1 for 24h. To analyse the cytotoxicity of MP-1, the spheroids were incubated with 5 µg/ml Hoechst 33342 for 20 min and further with 1.5 µg/ml PI for 5mins before analysing under the confocal microscope (Nikon A1R LSM). 405nm laser for Hoechst 33342 with excitation/emission wavelengths of 407nm/450nm and 488nm laser for PI with excitation/emission wavelengths of 488nm/525nm respectively was used to analyse the spheroids under the confocal microscope.

## Data availability

The data associated with this paper are openly available from the University of Leeds Data Repository. https://doi.org/10.5518/1558

## Author contributions

AB performed model membrane leakage assays, AFM studies and corresponding data analysis, contributed to manuscript writing. AP performed cell culture studies and corresponding data analysis, contributed to manuscript writing. DK performed model membrane leakage assays and corresponding data analysis. DD developed programming for the liquid handling robot, LK assisted with AFM studies and analysis. WB supervised patch clamp studies and contributed to writing. SDC: Project design, organisation and supervision, TH: Project design, organisation and supervision, manuscript writing, PAB: Project design, organisation and supervision, manuscript writing.

