## Supplementary Information for "Determinants of membrane sensitivity to the peptide MP-1 (*Polybia paulista*)"

Dr. Dagmara Kobza: Laboratory of Molecular Oncology and Innovative Therapies: Military Institute of Medicine - National Research Institute, Warsaw, Poland.

| | 1 | 2 | 3 | 4 | 5 | 6 | 7 | 8 | 9 | 10 | 11 | 12 | 13 | 14 |
|---|---|---|---|---|---|---|---|---|---|---|---|---|---|---|
| **MP-1** | **I** | **D** | **W** | **K** | **K** | **L** | **L** | **D** | **A** | **A** | **K** | **Q** | **I** | **L** |
| **IIA** | **A** | D | W | K | K | L | L | D | A | A | K | Q | I | L |
| **D2A** | I | **A** | W | K | K | L | L | D | A | A | K | Q | I | L |
| **W3A** | I | D | **A** | K | K | L | L | D | A | A | K | Q | I | L |
| **K4A** | I | D | W | **A** | K | L | L | D | A | A | K | Q | I | L |
| **K5A** | I | D | W | K | **A** | L | L | D | A | A | K | Q | I | L |
| **L6A** | I | D | W | K | K | **A** | L | D | A | A | K | Q | I | L |
| **L7A** | I | D | W | K | K | L | **A** | D | A | A | K | Q | I | L |
| **D8A** | I | D | W | K | K | L | L | **A** | A | A | K | Q | I | L |
| **K11A** | I | D | W | K | K | L | L | D | A | A | **A** | Q | I | L |
| **Q12A** | I | D | W | K | K | L | L | D | A | A | K | **A** | I | L |
| **I13A** | I | D | W | K | K | L | L | D | A | A | K | Q | **A** | L |
| **L14A** | I | D | W | K | K | L | L | D | A | A | K | Q | I | **A** |
| **A9Q** | I | D | W | K | K | L | L | D | **Q** | A | K | Q | I | L |
| **L7K** | I | D | W | K | K | L | **K** | D | A | A | K | Q | I | L |
| **A9K** | I | D | W | K | K | L | L | D | **K** | A | K | Q | I | L |
| **Q12K** | I | D | W | K | K | L | L | D | A | A | K | **K** | I | L |
| **I13K** | I | D | W | K | K | L | L | D | A | A | K | Q | **K** | L |
| **K4H** | I | D | W | **H** | K | L | L | D | A | A | K | Q | I | L |
| **K5H** | I | D | W | K | **H** | L | L | D | A | A | K | Q | I | L |
| **K11H** | I | D | W | K | K | L | L | D | A | A | **H** | Q | I | L |

**Figure S1:** Table of peptide variants and amino acid sequences. Red- negatively charged aspartic acid, green – positively charged lysine or histidine, blue - glutamine

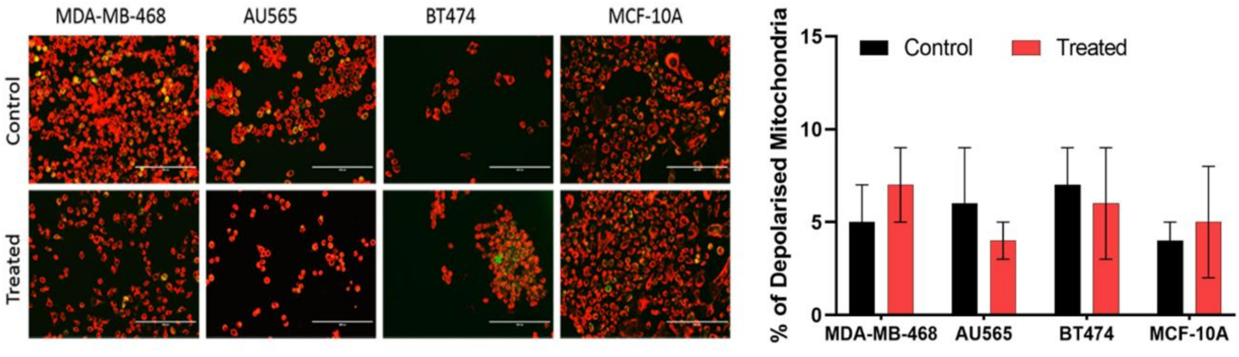

**Figure S2:** Mitochondrial membrane potential assay. Potential-driven accumulation of the JC-1 dye at mitochondrial membranes was quantified using fluorescence confocal microscopy to assess the impact of $ID_{50}$ MP-1 on mitochondrial membranes.

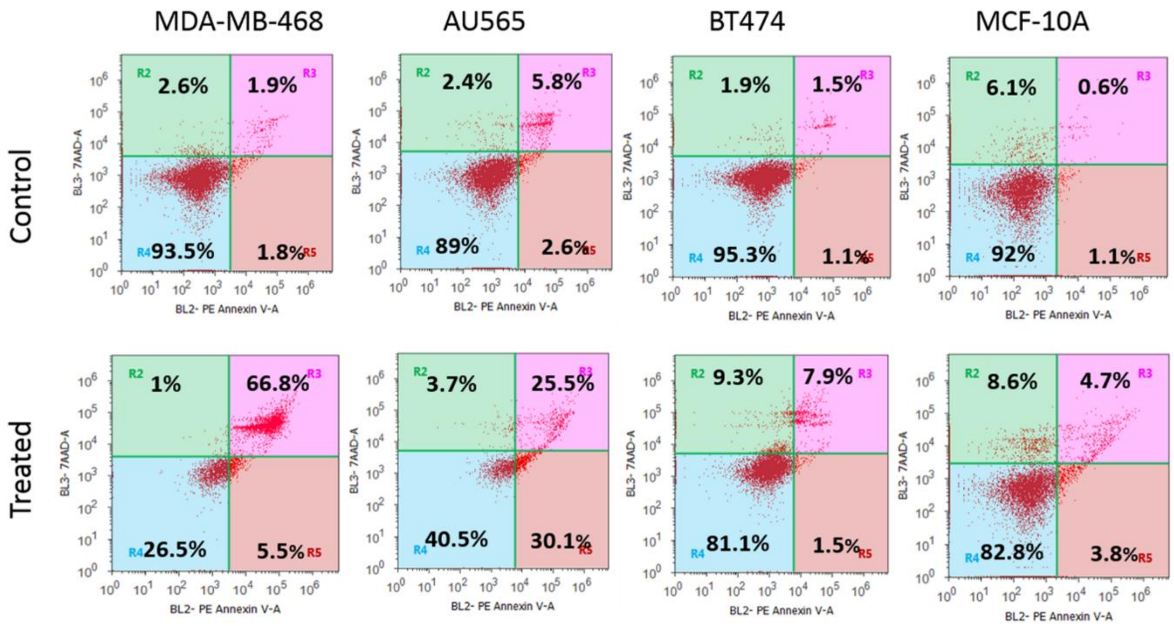

**Figure S3:** Flow cytometry data for MP-1 induced apoptosis assay (see Figure 1 E).

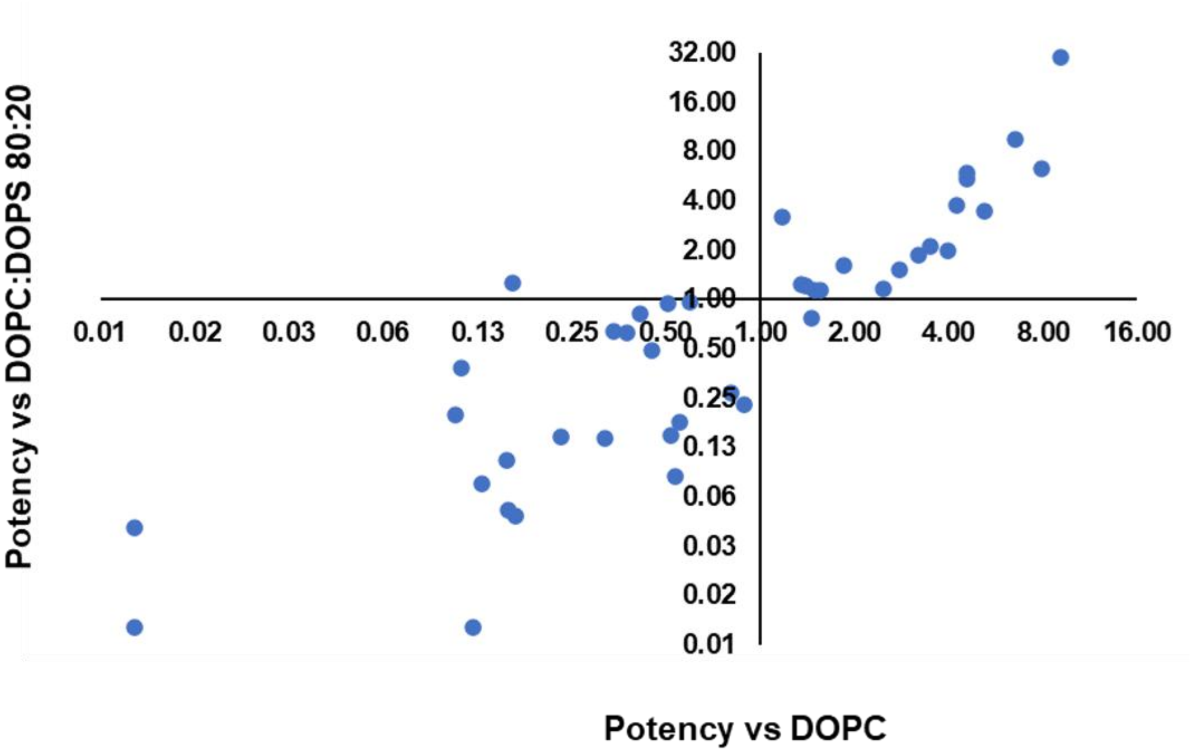

**Figure S4**: Log2 plot of potency relative to MP-1 for peptide variants against DOPC and DOPC:DOPS: 80:20 membranes. Pearson correlation value: 0.78464, p = <0.0001.

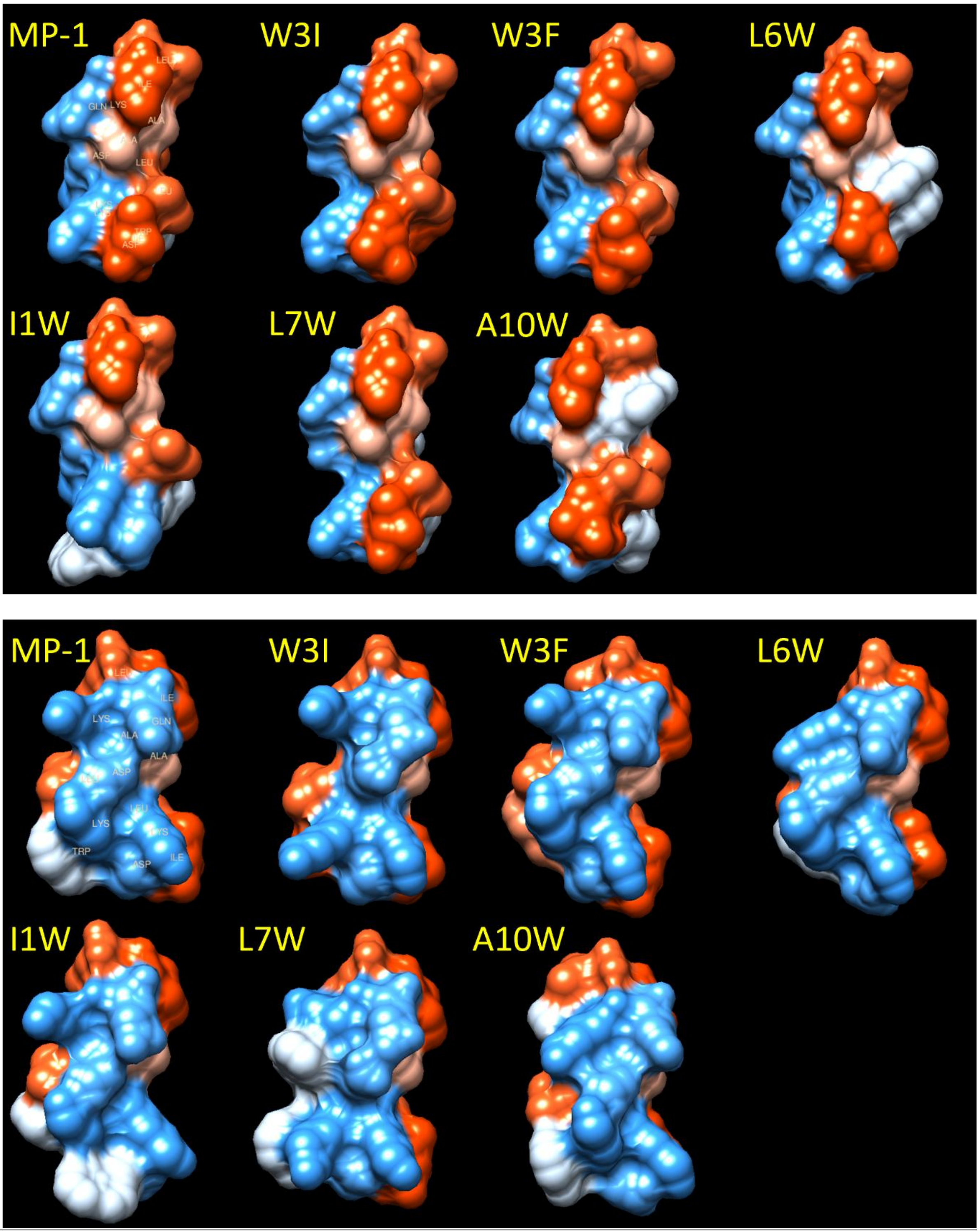

**Figure S5:** Kyte-Doolittle hydrophobicity surface space filling models MP-1 alongside W variants: red- hydrophobic, blue- hydrophilic, two projections are shown for each molecule, corresponding to a 180 degree rotation around the axis of the amide backbone, based on *de novo* structure prediction using the PEP-FOLD3 software[1], surface visualised using USCF Chimera[2].

| | Potency values | | IC$_{50}$ values ( μM) | | | |
|---|---|---|---|---|---|---|
| | 80:20 DOPC:DOPS | PC | 468 | 565 | 474 | 10A |
| **MP-1** | **1.00** | **1.00** | 23.4 | 18.6 | 50.61 | 233.6 |
| **D2K** | 9.21 | 4.25 | 16.81 | 33 | 88.5 | 140 |
| **D8K** | 47.93 | 6.30 | 13.56 | 23.1 | 51.76 | 55.23 |
| **D2A** | 0.72 | 0.94 | 11.74 | 13.52 | 28.73 | 26.25 |
| **L6W** | 5.04 | 0.81 | 73.26 | 100.2 | 144.7 | 155.1 |
| **I1A** | 0.15 | 0.30 | 102.2 | 428.5 | 207.2 | 157.6 |
| **L7K** | 0.02 | 0.00 | 104 | 87 | 416 | 214 |
| **D8A** | 1.62 | 3.68 | 9.82 | 12.55 | 26.23 | 20.03 |
| **A9Q** | 0.23 | 0.10 | 117.6 | 52.08 | 163.2 | 72.58 |
| **Q12A** | 1.55 | 0.24 | 18.47 | 11.34 | 28.11 | 67.43 |
| **I13K** | 0.06 | 0.21 | 1085 | 250 | 287 | 872 |
| **K4H** | 1.23 | 0.53 | 76.27 | 58.66 | 131.5 | 44.15 |
| **K11A** | 1.02 | 1.11 | 35.38 | 34.82 | 79.63 | 68.91 |

**Figure S6**: Table of peptides and corresponding potency values ($Potency = \frac{1}{\left(\frac{L_{50}Mutant}{L_{50}MP1}\right)}$) and IC$_{50}$ values.

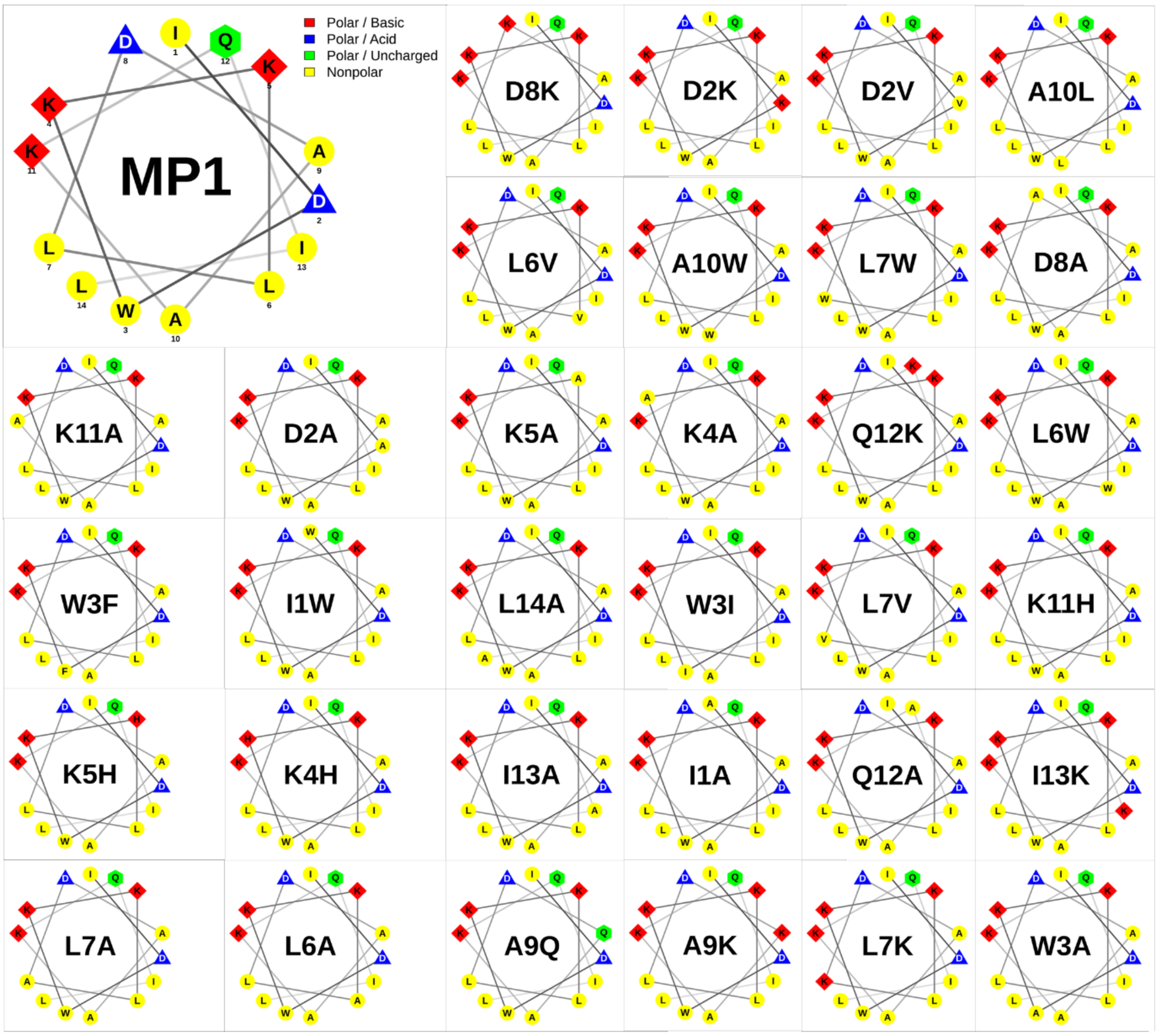

**Figure S7:** Helical wheel representations of the MP1 peptide and the derivatives investigated in this work. Projections were generated using the online NetWheels tool (http://lbqp.unb.br/NetWheels/).

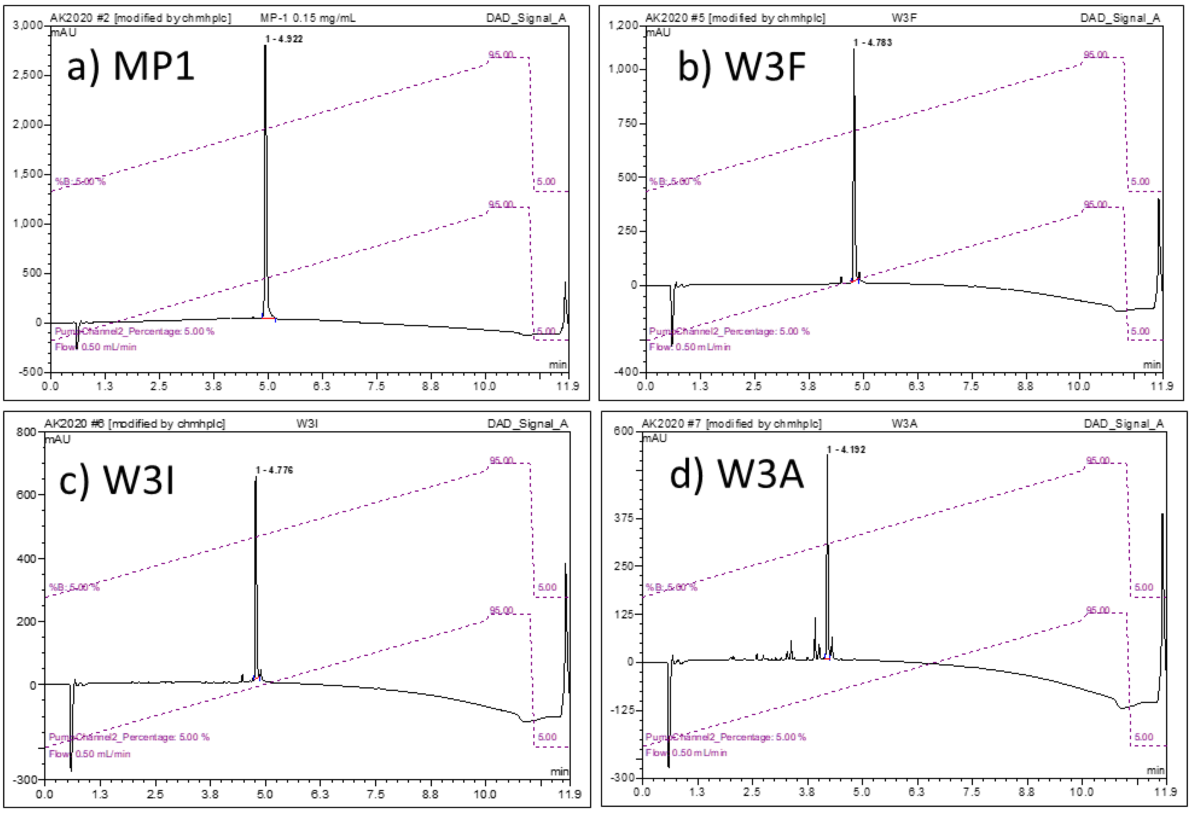

**Figure S8:** Analytical HPLC chromatograms of selected peptides.
